# Non-occluding Functions of Septate Junction Proteins in Signaling Regulation and Cell Polarity during Epithelial Development

**DOI:** 10.1101/2024.09.06.611635

**Authors:** Vanessa Holtwick, Hannah Müller, Andrea Schubert, Katja Rust

## Abstract

Occluding junctions are essential for epithelial tissue integrity and barrier function but also exert non-occluding roles. Here we identify a several crucial function of the invertebrate occluding junction components Coracle, Nervana 2, Neurexin-IV, and Kune-kune in regulating cell polarity during follicle epithelium development. We show that the morphogenesis of the follicular stalk is an actin-driven process that requires intact apical-basal cell polarity, which is controlled by septate junction components. Occluding junction components further regulate signaling pathways in a cell-type specific manner. In undifferentiated stem cells and immediate daughters, septate junction components promote effective Wnt signaling to control proliferation, while they limit Jak-STAT signaling activity induced by polar cells. Together, our data emphasize the multiple roles of occluding junction components independent of their classical role in forming the paracellular barrier.

## Introduction

Functional epithelial tissues are vital for organismal development and health. Epithelial homeostasis and development require a tight coordination of proliferation, differentiation and cell morphological changes. The *Drosophila* follicle cell lineage depicts an excellent model to study epithelial development from a highly proliferative niche-maintained stem cell to a single layered mature epithelium (Fig. 1A). Follicle cells are produced in the germarium by follicle stem cells (FSCs), which obtain niche signals from neighboring escort cells and cap cells (Forbes *et al*, 1996; Sahai-Hernandez & Nystul, 2013; Song & Xie, 2003; Wang & Page-McCaw, 2014). FSCs divide to produce transiently amplifying prefollicle cells (pFCs) which associate with developing germline cysts and differentiate. The first differentiation choice pFCs can undergo is induced by Delta, which is produced by germline cysts in Region 2b (Assa-Kunik *et al*, 2007; Dai *et al*, 2017; Rust *et al*, 2020). Delta receiving pFCs differentiate into polar cells, which form two clusters of signal producing cells on either side of the germline cyst. Polar cells then produce the Jak-STAT ligand Unpaired (Upd) and pattern the surrounding follicle epithelium. Neighboring follicle cells that receive Upd in the germarium differentiate into stalk cells and form a single row of cells connecting two neighboring germline cysts (Assa-Kunik *et al*, 2007; McGregor *et al*, 2002; Rust & Nystul, 2020). The remaining pFCs differentiate into main body follicle cells, which organize into a single layer epithelium that ensheaths the germline cyst. (Fig. 1A).

**Figure 1:**
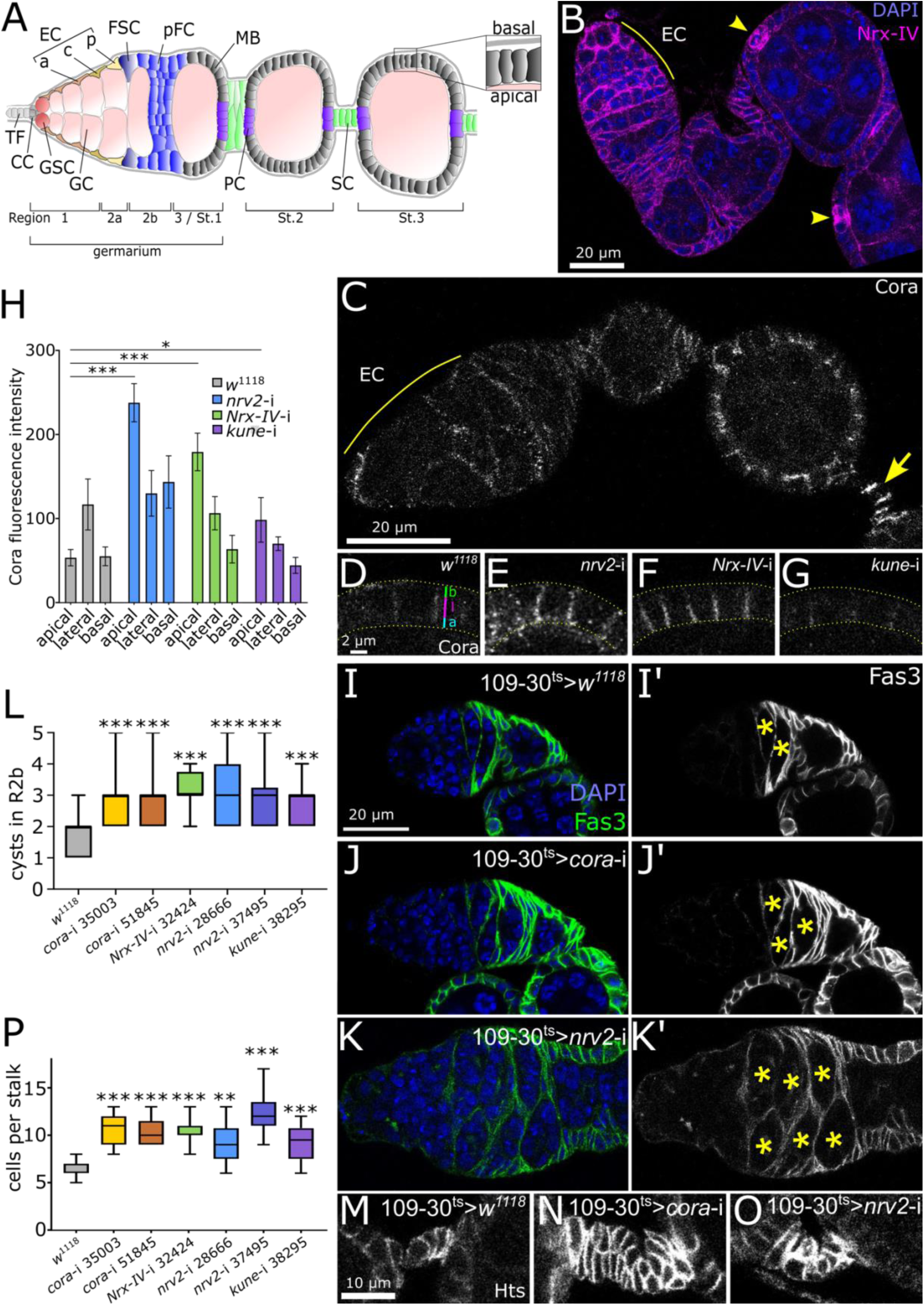
SJ proteins regulate early follicle epithelium development. A) Graphic representation of the *Drosophila* ovary and the early stages of oogenesis. TF: terminal filament, CC: cap cell, GSC: germline stem cell, GC: germ cell, EC: escort cell, a: anterior, c: central, p: posterior, FSC: follicle stem cell, pFC: prefollicle cell, MB: main body follicle cell, PC: polar cell, SC: stalk cell. Inset on the right highlights the apical to basal orientation of main body follicle cells. B) Germarium and early cyst stages stained for Nrx-IV::mCherry (magenta) and DAPI (blue). Nrx-IV::mCherry is highly expressed in escort cells (EC) and polar cells (arrowheads). C) Wildtype ovariole (109-30^ts^ > *w^1118^*) stained for Cora. Cora is expressed in escort cells and follicle cells with high expression in mature stalk (arrow). D-H) Immunostaining (D-G) and associated quantification (H) of Cora staining in Stage 6 main body follicle. 109-30^ts^ was used to induce the respective RNAi. In wildtype Cora mostly localizes laterally. Knockdown of *nrv2*, *Nrx-IV* or *kune* disturbs normal Cora localization. Label in D) illustrates how we divided the lateral membrane into the apical (“a”), lateral (“l”) and basal (“b”) portions. H) n = 6, 7, 7, 6 measurements for each condition. I-K) Germaria of the indicated genotype stained for DAPI (blue) and the follicle cell marker Fas3 (green or white in I’-K’). Germline cysts encapsulated by follicle cells in the germarium are marked by asterisks. L) Quantification of cysts in Region 2b in 109-30^ts^ crossed with the respective genotype. n = 37, 22, 21, 36, 42, 14, 15 germaria respectively. M-O) Stalk regions of the genotypes indicated in each respective row stained for Hts. Note that wildtype stalk forms a single row of cells (M) while knockdown of SJ components results in double rowed stalks (N-O). P) Quantification of cells per stalk (Stage 4) in ovarioles from 109-30^ts^ crossed to the respective allele. n = 28, 11, 9, 12, 6, 28, 12 stalks respectively.

The follicle epithelium possesses classical epithelial properties, including apical-basal polarity and cellular junctions (Franz & Riechmann, 2010; Müller, 2000). Among epithelial junctions, septate junctions (SJs) are invertebrate occluding junctions that localize to the lateral membrane and assume a barrier function similar to vertebrate tight junctions (Izumi & Furuse, 2014). Molecularly, SJs are conserved to vertebrate SJs, known as axo-glial SJs, in myelinated axons (Banerjee *et al*, 2006). Based on their appearance in ultrastructural microscopy and molecular composition, mature SJs are classified in either pleated, ladder-like SJs (pSJ) or smooth SJs (sSJ). Further, SJs can form as tricellular junctions (tSJ). SJs are formed by roughly thirty different components out of which seventeen have been identified as core components (Rice *et al*, 2021). Studies using the *Drosophila* embryonic epithelium have identified several SJ core components as critical cell polarity regulators, where this group of proteins maintains basolateral membrane identity by restricting the apical polarity protein Crumbs (Laprise *et al*, 2009). Among these are the transmembrane proteins Neuroglian (Nrg) and Neurexin-IV (Nrx-IV), the FERM domain protein Coracle (Cora), as well as the α and β subunits of the Na^+^/K^+^ ATPase ATPα and Nervana 2 (Nrv2), which both assume ion-pump-independent roles in SJ formation. However, it is unclear whether all SJ core components, including the Claudin homolog Kune-kune (kune), are required for cell polarity. Moreover, SJ components fulfill further non-occluding functions in other systems, where they regulate processes like cell migration, cell-cell communication, cell adhesion and proliferation (Rice *et al*, 2021; Rouka *et al*, 2021). In the follicle epithelium, SJ components are universally expressed (Alhadyian *et al*, 2021). Yet, SJs only assemble into incipient SJs at Stage 6 and are visible as mature, pleated SJs from Stage 10 onwards (Müller, 2000), suggesting that SJ components assume non-occluding functions during follicle epithelium development. In the late follicle epithelium, SJ proteins are required for egg elongation and the formation of dorsal appendages, which are formed from late stage main body follicle cells (Alhadyian *et al*, 2021). They are also required for border cell migration, where the anterior polar cells and surrounding outer border cells delaminate and migrate through the nurse cells towards the oocyte (Alhadyian *et al*, 2021; Felix *et al*, 2015). However, the mechanism behind SJ component function in follicle epithelium morphogenesis is unclear.

Here we study the SJ core components Cora, Kune, Nrv2 and Nrx-IV in the early follicle cell lineage of *Drosophila melanogaster*. We identify several cell type specific and non-occluding roles of SJ components, which are crucial for normal follicle cell development. Our study indicates that SJ components are crucial for normal apical-basal polarity of the follicle epithelium, thereby regulating a range of processes. We show, for the first time, that SJ components regulate Wnt signaling activity in FSCs and pFCs. This grants normal proliferation and ensures the encapsulation of germline cysts. Further, we characterize the morphogenesis of the follicular stalk and show that this process is dependent on E-Cadherin, apical-basal polarity and the actin cytoskeleton, all of which are coordinated by SJ components. Lastly, SJ components are required in polar cells, where they regulate cell polarity and orient the microtubule network. This is required for apical transport of the mRNA encoding Unpaired (Upd), a JAK-STAT ligand, which is transported by the RNA-binding protein Cabeza (caz). Loss of SJ components disturbs the apical enrichment of Upd protein, thereby increasing JAK-stat activity in the basally located stalk cells, promoting stalk cell differentiation and resulting in supernumerary stalk cells.

## Results

### Normal follicle epithelium formation relies on SJ genes

SJ components are highly expressed in differentiated main body follicle cells (Alhadyian *et al*, 2021) but their expression during early follicle epithelium formation is poorly described. Using our single cell RNA-sequencing dataset of the *Drosophila* ovary (Rust *et al*, 2020), we investigated the expression of genes encoding SJ components in somatic cells of early stages of oogenesis. SJ junction genes including *cora* and *nrv2* display high expression in escort cells (Fig. S1A). Polar cells exhibit enriched expression for many SJ genes including *Gli*, *bark* and *M6*, which encode components contributing to tricellular SJs. *Nrg* is further highly expressed in stalk cells, while other follicle cell types express lower levels of SJ genes. Indeed, Nrx-IV-mCherry is expressed in all somatic cells of the ovary with higher levels in escort cells and strongest expression in polar cells, as predicted (Fig. 1B). Further, using antibody staining we detected Cora expression in escort cells as well as all types of follicle cells (Fig. 1C). In follicle cells Cora localized to the lateral membrane and we detected the highest Cora staining in mature stalk cells.

To investigate the function of SJs in the follicle epithelium we examined knockdown of the core SJ components *cora*, *nrv2*, *Nrx-IV* and *kune*. To this aim, we employed the pan-follicle cell Gal4 driver 109-30-Gal4 (Hartman *et al*, 2015) combined with tub-Gal80^ts^ (hereafter referred to as ^ts^) to target the adult follicle epithelium (Fig. S1B). To validate the knockdown of *cora* we stained the ovaries with *cora* knockdown in follicle cells with an anti-Cora antibody. When compared to control, Cora staining was strongly reduced in follicle cells while escort cells, which are not targeted by 109-30^ts^, displayed unaltered Cora expression (Fig. S1C-D). SJ components including Cora localize laterally (Tiklová *et al*, 2010). Consistently, Cora staining was highest in the lateral domain of wildtype main body cell membranes in Stage 6 (Fig. 1D). In contrast, knockdown of *kune*, *nrv2* or *Nrx-IV* shifted Cora localization towards the apical portion of the lateral cell membrane (Fig. 1E-H), in agreement with the interdependent localization of SJ components.

Having confirmed the functionality of RNAi lines, we next examined the morphological phenotypes of ovaries upon 109-30^ts^-induced knockdown of *cora*, *kune*, *nrv2* or *Nrx-IV*. Interestingly, we found that depletion of *cora*, *kune*, *nrv2* or *Nrx-IV* resulted in similar morphological defects. While wildtype germaria contained on average 1.76 ± 0.1 follicle cell encapsulated cysts in Region 2b, knockdown of these SJ proteins significantly increased the number of follicle cell encapsulated cysts in this region (Fig. 1I-L, S1E-F, *cora* #35003: 2.86 ± 0.19, *cora* #51845: 2.86 ± 0.19, *kune* #38295: 2.67 ± 0.16, *nrv2* #28666: 3.07 ± 0.14, *nrv2* #37495: 2.93 ± 0.25, *Nrx-IV* #32424: 3.06 ± 0.11 cysts per Region 2b). Only 6.47 % of control germaria contained more than two cysts in Regions 2b, while knockdown of SJ components significantly increased the percentage of ovaries with more than two cysts (Fig. S1G). Further, when we stained ovarioles for Hu li tai shao (Hts), which is expressed in the entire follicle epithelium, we noted that stalks were double rowed in comparison to the single row morphology observed in wildtype (Fig. 1M-O, S1H-I). While wildtype stalks contained on average 6.86 ± 0.15 cells per stalk, knockdown of SJ components *cora*, *kune*, *nrv2* or *Nrx-IV* significantly increased stalk cell numbers (Fig. 1P). The morphological defects caused by depletion of *cora*, *nrv2*, *Nrx-IV* or *kune* indicate that these SJ proteins are vital for follicle cell morphogenesis and encouraged us to further explore their function in follicle epithelial development.

### SJ components regulate proliferation of undifferentiated follicle cells via Wnt signaling

Loss of SJs affects proliferation in other types of epithelia (Chen *et al*, 2020; Izumi *et al*, 2019; Ward *et al*, 2001). Thus, the expansion of the stalk cell number upon SJ gene knockdown prompted us to investigate proliferation. We quantified the percentage of EdU labeled cells upon pan-follicle knockdown of SJ genes and found that, as in the wildtype, stalk cells are quiescent outside of the germarium (Fig. S3A-C, no EdU positive stalk cells from 109-30^ts^ crossed to *w^1118^*, *cora* #35003, *cora* #51845, *kune* #38925, *nrv2* #28666, *nrv2* #37495, *Nrx-IV* #32424, n = 9, 11,10, 14, 10, 11, 8 ovarioles respectively). The expansion of the stalk cell pool is therefore not caused by a failure of differentiated stalk cells to exit mitosis. To investigate whether the increased stalk cell numbers might be caused by an elevated proliferation rate of FSCs or pFCs we quantified EdU positive somatic cells in Region 2b of the germarium. We found that knockdown of SJ genes with the pan-follicle cell driver 109-30^ts^ significantly reduced the percentage of EdU positive follicle cells in Region 2b (Fig. 2A, *w^1118^*: 22.9 ± 3.27%, *cora* #35003: 6.55 ± 0.93%, *cora* #51845: 11.26 ± 2.81%, *kune* #38295: 8.73 ± 1.91%, *nrv2* #28666: 12.18 ± 2.64%, *nrv2* #37495: 7.71 ± 1.99%, *Nrx-IV* #32424: 9.42 ± 3.05%). To test whether SJ component knockdown generally reduced proliferation of dividing cells, we assessed EdU incorporation of main body follicle cells shortly before they exit mitosis in Stage 6. However, EdU incorporation in main body follicle cells was not affected (Fig. 2B).

**Figure 2:**
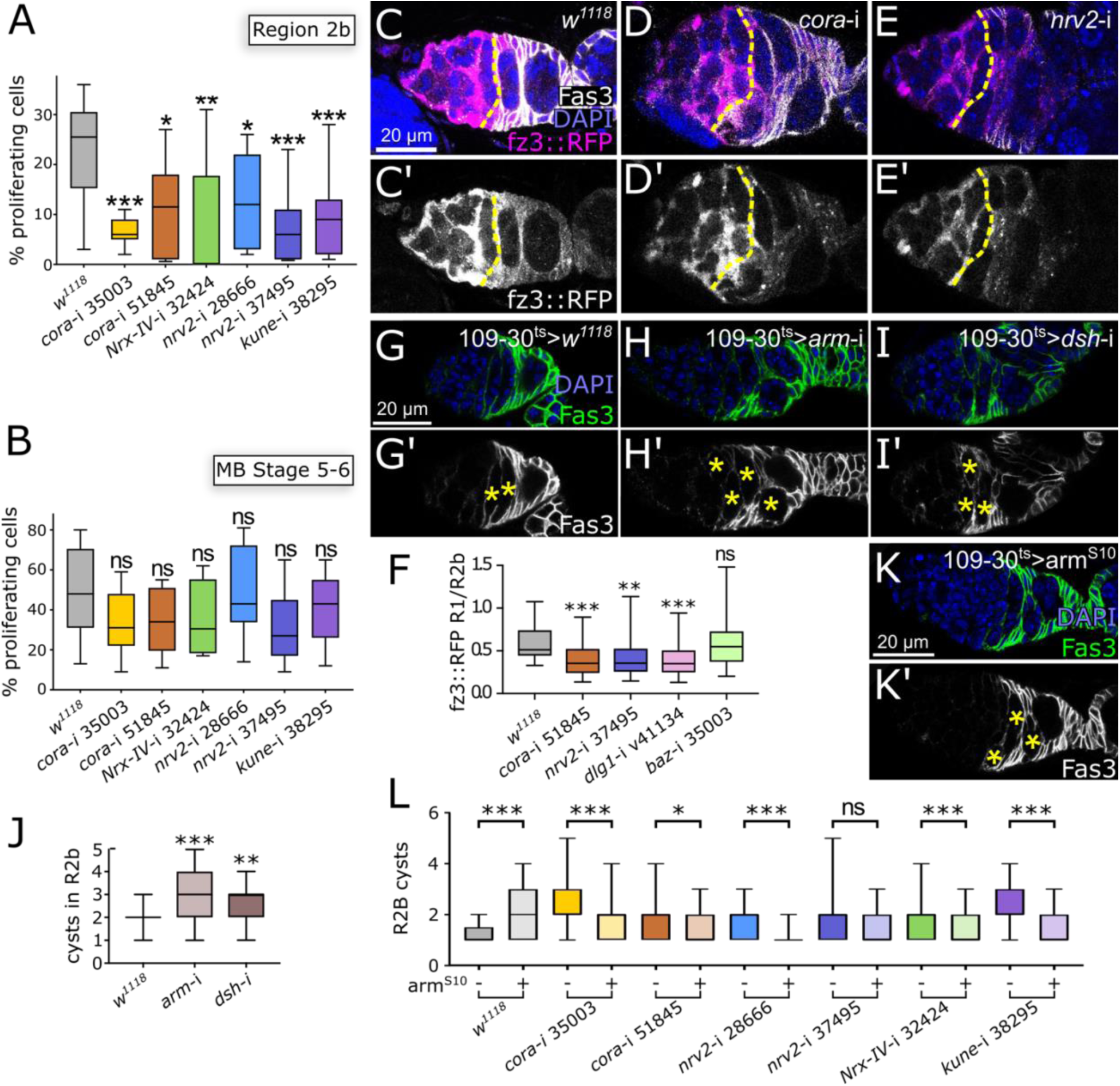
SJ proteins promote Wnt signaling to regulate proliferation in early follicle cells. A-B) Quantification of the percentage of proliferating cells in Region 2b (A) or main body follicle cells of Stage 5-6 cysts (B) in ovarioles of 109-30^ts^ crossed to the respective allele. A) n = 10, 11, 10, 12, 11, 11, 15 germaria. B) n ≥ 9, 11, 9, 8, 10, 11, 13 ovarioles. C-E) Germaria of 109-30^ts^ crossed to fz3::RFP and the respective allele and stained for DAPI (blue), RFP (magenta) and Fas3 (white). C’-E’) show the RFP channel in white. Yellow dotted line shows the Fas3 border between Region 2a and 2b. F) F) Quantification of fz3::RFP intensity. To ensure robustness, we normalized measurements in Region 2b (affected by Gal4) to measurements in Region 1 ECs (not affected by Gal4). Control (*w^1118^*) and RNAi-lines were induced with 109-30^ts^. n = 40, 40, 30, 31, 40 germaria respectively. G-I) Ovarioles of the 109-30^ts^ driver crossed to the wildtype *w^1118^*(G), *arm*-RNAi (H) or *dsh*-RNAi (I) and stained for DAPI (blue) and Fas3 (green). G’-I’) show Fas3 in white. Asterisks mark Region 2b cysts. J) Quantification of cysts in Region 2b in the wildtype (109-30^ts^ > *w^1118^*) or upon reduction of Wnt signaling by RNAi depletion of *arm* or *dsh* in follicle cells with 109-30^ts^. n = 26, 21, 18 germaria respectively. K) Germarium with constitutively active Wnt signaling induced by arm^S10^ stained for DAPI and Fas3. Region 2b cysts are marked with yellow asterisk. L) Quantification of Region 2b cysts in 109-30^ts^ inducing the respective genotype and in the presence or absence of arm^S10^. n = 57, 90, 55, 84, 58, 75, 60, 74, 56, 59, 71, 77, 54, 73.

Various signaling pathways regulate proliferation in undifferentiated follicle cells, including the Wnt signaling pathway, which is active in FSCs (Rust and Nystul, 2020; Song and Xie, 2003). We tested the influence of *cora* and *nrv2* knockdown on the Wnt pathway reporter fz3::RFP and found a significant reduction of Wnt signaling activity in Region 2b follicle cells (Fig. 2 C-E, S2D). Therefore, SJ proteins likely regulate proliferation of undifferentiated follicle cells via promoting Wnt pathway activity (Fig. 2A). As SJs are maturely formed in early follicle epithelium development (Müller, 2000), we assumed that SJ proteins regulate Wnt signaling independent of barrier formation. One prominent non-occluding function of Cora, Nrv2 and Nrx-IV is the regulation of cell polarity (Laprise *et al*, 2009). Therefore, we tested whether manipulation of classical cell polarity regulators also affects Wnt signaling activity. We reanalyzed the effects of SJ protein depletion on fz3::RFP this time normalizing each data point, confirming that cora or nrv2 loss significantly reduced fz3::RFP intensity. (Fig. 2F). Depletion of *discs large-1 (dlg1)*, which encodes an important basolateral regulator of cell polarity, similarly significantly reduced fz3::RFP intensity (Fig. 2F, S2E-F). In contrast, depletion of *bazooka (baz, Drosophila* Par-3), encoding an apical polarity protein, did not affect fz3::RFP levels (Fig. 2F, S2E,G). This agrees with a previous report showing that Dlg-1 but not Baz display polarized localization in FSCs (Kronen *et al*, 2014; Franz & Riechmann, 2010). Next, we tested whether genetic reduction of Wnt signaling activity by RNAi-mediated depletion of *Drosophila* β-Catenin *armadillo* (*arm*) or *dishevelled* (*dsh*), encoding a Wnt pathway component, phenocopied SJ component depletion. Indeed, reduction of Wnt signaling by RNAi-mediated depletion of the genes encoding the Wnt signaling effector β-Catenin (*arm*) or the positive regulator of Wnt signaling *dishevelled* (*dsh*) resulted in an increase of germline cysts in Region 2b, similar to the knockdown of SJ core components (Fig. 2G-J, *w^1118^*: 1.88 ± 0.08, *arm*: 3.38 ± 0.21, *dsh*: 2.56 ± 0.23 cysts per germarium). Lastly, we tested whether Wnt pathway activation by expression of the constitutively active β-Catenin construct arm^S10^ can rescue cyst accumulation upon SJ protein depletion. Surprisingly, germaria with follicle cell specific expression of arm^S10^ showed supernumerary cysts in Region 2b (Fig. 2K-L). This indicates that the balance of follicle cells and germline cells needs to be tightly regulated to efficiently shuttle germline cysts through the germarium. In agreement, arm^S10^ largely restored Region 2b cyst numbers in the background of SJ component depletion (Fig. 2L). Together, SJ components and Dlg1, another cell polarity regulator, promote Wnt pathway activity to ensure sufficient proliferation of undifferentiated follicle cells. This is crucial for efficient germline cyst shuttling through the germarium. Further, the double row stalk cell phenotype observed upon SJ gene depletion is not caused by excessive proliferation of FSCs or pFCs.

### SJ components regulate actin and cell polarity to promote stalk morphogenesis

In order to examine whether the double row stalk phenotype is caused by faulty stalk cell differentiation, we examined the stalk cell markers Anterior open (Aop) and Laminin C (LamC), Eyes absent (Eya), which is absent during normal stalk cell development, and Zfh1, which is expressed in undifferentiated follicle cells and retained during stalk cell differentiation (Boisclair Lachance *et al*, 2014; Pearson *et al*, 2016; Rust *et al*, 2020; Bai & Montell, 2002). Examination of ovaries with knockdown of SJ genes revealed that neither of these cell fate markers were misexpressed, suggesting that stalk cells differentiate normally (Fig. S4A-T). A previous study claimed that stalk cell numbers are fine-tuned via stalk cell apoptosis (Borensztejn *et al*, 2018). However, while staining for the apoptosis marker Dcp-1 successfully identified polar cell death (Fig. S4U), we never observed apoptotic stalk cells. Next, we sought to investigate whether SJ components directly affect stalk development. Normal stalk morphogenesis relies on a process called cell intercalation, where cells from different layers integrate into a single row of cells. SJ components regulate cytoskeletal organization during cell intercalation in embryonic dorsal closure (De *et al*, 2022). Since the importance of cell polarity and cytoskeleton dynamics has not been investigated during stalk development, we first sought to characterize stalk morphogenesis in more detail. Previously, stalk development has been classified into distinct steps starting: Initiation in Region 2b, where stalk cells are specified; Convergence in Region 3, where stalk cells start contacting each other; Intercalation into a single row stalk in Stage 3; and Closing in Stages 4-5 where the extracellular matrix is reorganized (Van De Bor *et al*, 2021). We stained wildtype stalk for E-Cadherin (E-Cad), as a marker for adherens junctions, and Phalloidin to visualize the F-actin cytoskeleton (Fig. 3A-F). During initiation E-Cad and F-actin start to accumulate between stalk cell precursors located between the two lens-shaped germline cyst in Regions 2b and 3 (Fig. 3A, G). When germline cysts are advanced in Region 3, characterized by rounded cysts, E-Cad and F-actin are accumulated between stalk cells, which have started constricting (Fig. 3B, G). As cysts advance further they bud off the germarium, which is characterized by a morphologically recognizable stalk region. Here stalk cells have formed a double row and display a consistent line of E-Cad and high F-actin between them, which forms a “zipper” line across the intercalating stalk cells (Fig. 3C, G). Intercalation does not occur between all stalk cells simultaneously. Instead, the first stalk usually displays mostly a single row of stalk cells while few stalk cells are still in the process of intercalation (Fig. 3D, G). The second stalk is often fully intercalated, although we noticed some variation, and here stalk cells display spots of E-Cad between them (Fig. 3E, G). The third stalk is normally matured and displays E-Cad localization laterally between stalk cells (Fig. 3G-G). To better understand the intercalation process, we applied live imaging using endogenously tagged E-Cad. We found that the process from budding, where a “zipper” line of E-cad is visible between a double row of stalk cells, to intercalation, where E-Cad is mostly visible as spot-like accumulations between stalk cells, occurs in a matter of a few hours (Fig. 3H-J, Supplementary Movie 1).

**Figure 3:**
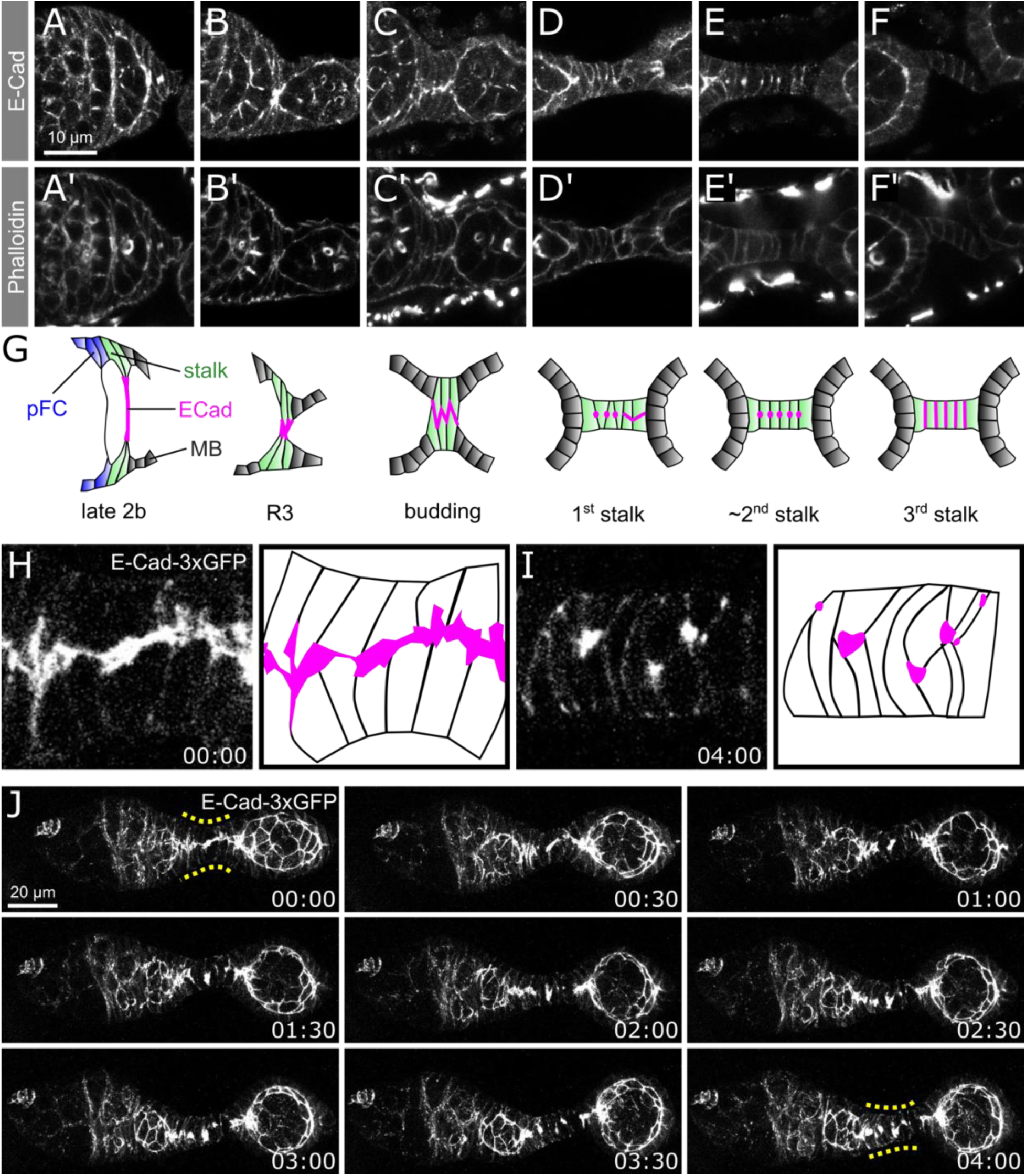
Distinct steps of stalk cell intercalation. A-F) Consecutive stages of stalk morphogenesis in wildtype ovaries of the genotype 109-30^ts^ > *w^1118^* stained for E-Cad (A-F) and Phalloidin (A’-F’). A graphic representation of stalk morphogenesis is depicted in G). h-j) Snapshots of live imaging analysis of E-Cad-3xGFP during stalk morphogenesis from the budding stage (H) to the first stalk (I). J) Snapshots of a maximum intensity projection in 30 minute intervals. For video see supplementary movie 1. Yellow dotted line marks the stalk region.

As the double row phenotype evoked by knockdown of *cora*, *nrv2*, *Nrx-IV* and *kune* indicates defects in stalk cell intercalation, we focused on the budding stage, where stalk cells are present in a double row in the wildtype (Fig. 1M-O, S1G-H, 3C). Using three-dimensional image analysis, we confirmed that E-Cad localizes in a single, consistent line between stalk cells in the wildtype (Fig. 4A, note the cross-sections). In contrast, pan-follicle cell knockdown of *cora*, *nrv2*, *Nrx-IV* and *kune* resulted in a range of defects including the formation of multiple E-Cad lines, indicating incomplete contact between stalk cells, inconsistent and broken E-Cad lines and mislocalization to the outer, basal side of stalk cells (Fig. 4B-C, S5A-B).

**Figure 4:**
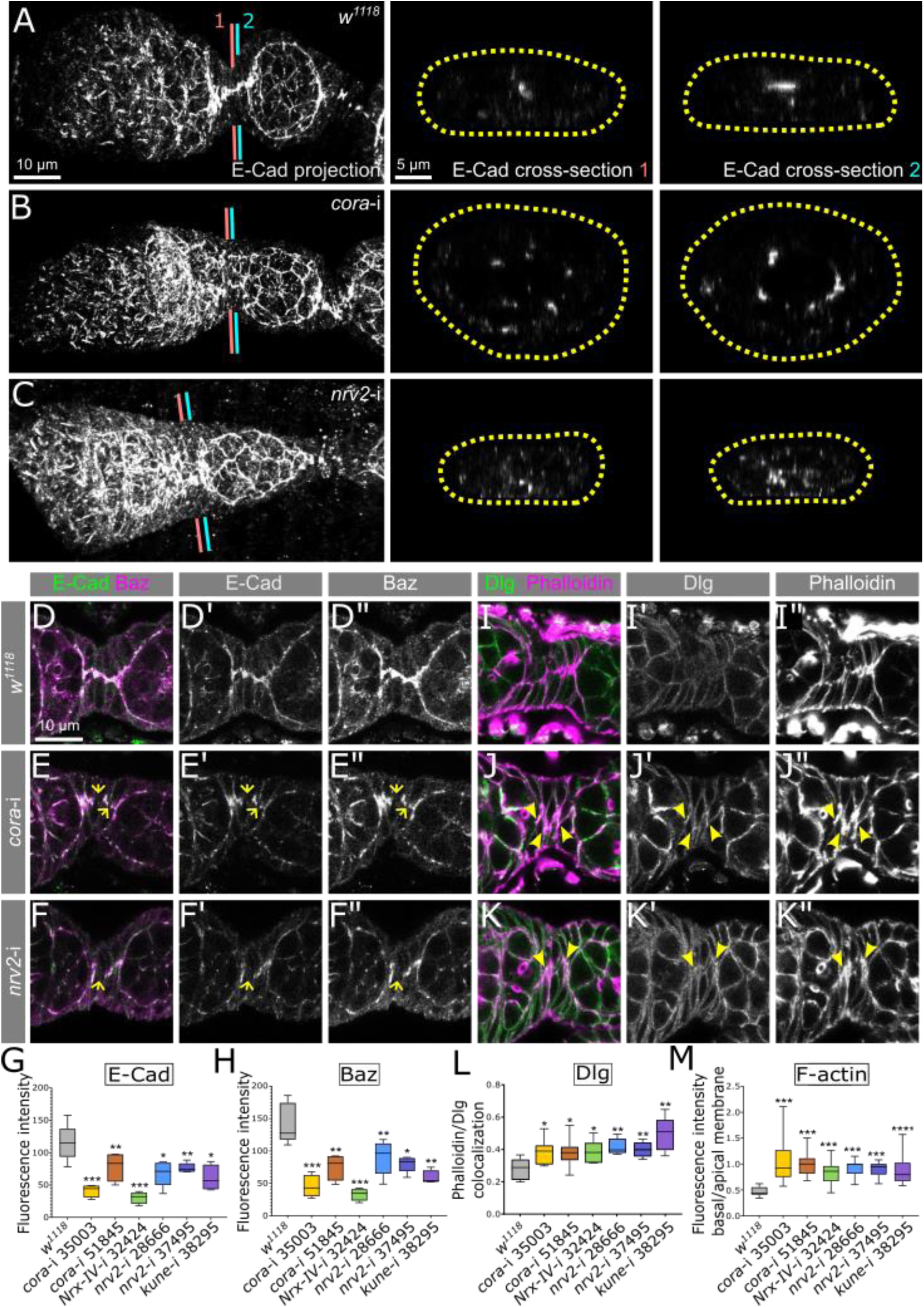
SJ components regulate apico-basal polarity to govern stalk cell intercalation. A-C) Imaris analysis of E-Cad expression in ovarioles from 109-30^ts^ crossed to the indicated genotype. Cross-sections (right) are labeled in the maximum intensity overview (left). Yellow dotted line outlines the stalk diameter. D-G) Budding stalk regions of ovarioles from 109-30^ts^ crossed to the allele indicated in each row stained for E-Cad (green) and Baz (magenta) (D-F) or E-Cad (D’-F’) or Baz (D”-F”) in white. Arrows indicate regions of apical membrane lacking E-Cad and Baz staining. G-H) Quantification of E-Cad (G) and Baz (H) staining across the zipper line in Region 3 intercalating stalks of 109-30^ts^ crossed to the respective allele. n = 5, 4, 4, 4, 5, 5, 4 ovarioles respectively for G) and H). I-K) Budding stalk regions of 109-30^ts^ crossed to the indicated genotype stained for Dlg (green) and Phalloidin (magenta) (I-K) or Dlg (I’-K’) or Phalloidin (I”-K”) in white. Arrowheads indicate regions with co-localization of Dlg and F-actin along the apical membrane. L) Pearson’s coefficient describing the colocalization of Dlg and F-actin along the apical zipper line. n = 6, 8, 7, 6, 7, 6, 6 respectively. M) Quantification of basal to apical Phalloidin staining in budding stalk cells in 109-30^ts^ crossed to the respective genotype. n = 11, 15, 9, 14, 18, 16, 17 ovarioles respectively.

Next, we assessed markers of apical-basal polarity. Baz localizes to the E-Cad zipper line in budding stalks in wildtype (Fig. 4D). Knockdown of SJ components resulted in inconsistent zipper lines for E-Cad as well as Baz (Fig. 4E-F, S5C-D). Additionally, E-Cad and Baz expression were reduced as indicated by a reduced fluorescence intensity upon knockdown of *cora*, *nrv2*, *Nrx-IV* or *kune* (Fig. 4G-H). Dlg-1 was absent from the zipper line in wildtype and instead localized to the lateral membranes of budding stalk cells (Fig. 4I). In contrast, upon SJ component knockdown, Dlg-1 frequently co-localized with F-actin at the zipper line (Fig. 4J-L, S5E-F). Lastly, wildtype budding stalk cells displayed high levels of F-actin at the apical zipper line and lower F-actin levels at the basal side, while budding stalk cells with knockdown of *cora*, *nrv2*, *Nrx-IV* or *kune* did not establish this gradient (Fig. 4M). To test whether these defects are indeed responsible for the double row stalk phenotype, we depleted *E-Cad* from follicle cells by RNAi-knockdown. Indeed, knockdown of *E-Cad* resulted in cyst fusions and double row stalks, indicating defects in stalk cell intercalation (Fig. S5G). In summary, SJ components regulate cell polarity, adhesion and F-actin localization to promote stalk cell intercalation.

### SJ components control stalk cell number via Jak-STAT signaling

While the defective actin cytoskeleton and cell polarity causes stalk cell intercalation resulting in double rowed stalks observed in SJ knockdown, it is unlikely that this causes the increased stalk cell number (Fig. 1P). Accordingly, stalk cell specific knockdown of SJ components with the stalk cell specific driver CG46339^ts^ (Rust *et al*, 2020) resulted in double rowed stalks but did not increase stalk cell numbers (Fig. 5A-B). Instead, we postulated that an increased number of stalk cells is specified from the pFC pool. Stalk cell specification is regulated by polar cells in the germarium, which produce the Unpaired-ligand (Upd) to induce Jak-STAT signaling in neighboring pFCs which then differentiate towards the stalk cell fate (McGregor *et al*, 2002). First, we examined polar cell numbers upon SJ component depletion in follicle cells, hypothesizing that an increased number of polar cells would induce more cells to specify as stalk cells. We found that similar to wildtype ovaries, ovaries with knockdown of SJ components contained on average two polar cells (wildtype: 1.87 ± 0.06 polar cells, cora #35003: 1.83 ± 0.11, cora #51845: 1.91 ± 0.09, kune #38295: 2 ± 0, nrv2 #28666: 1.96 ± 0.07, nrv2 #37495: 1.92 ± 0.1, Nrx-IV #32424: 1.93 ± 0.08 cells per anterior polar cell cluster in Stage 3-4, p = ns, n = 31, 12, 11, 7, 25, 24, 30 respectively). Next, to test whether faulty Upd distribution from polar cells causes elevated stalk cell numbers, we targeted SJ genes for knockdown with the polar cell driver upd^ts^ (Bai & Montell, 2002). Indeed, polar cell specific knockdown resulted in a significant expansion of stalk cell numbers and was associated with a double row phenotype (Fig. 5C-F). Thus, SJ components function non-cell autonomously to regulate stalk cell numbers via polar cells. Consistently, when we examined the Jak-STAT reporter 2xSTAT-GFP we found that knockdown of SJ components significantly increased Jak-STAT signaling levels in stalk cells and main body follicle cells, in which Jak-STAT signaling is induced by polar cells (Fig. 5 G-K). In contrast, Jak-STAT signaling was not elevated in the FSC/pFC Region 2b, where Jak-STAT signaling is induced independently of polar cells (Fig. 5J). Hence, the increased stalk cell number upon SJ gene knockdown is caused by elevated Jak-STAT signaling induced by polar cells. We scrutinized whether polar cell polarity may regulate Jak-STAT activity in stalk cells. Polar cell specific depletion of the apical proteins Baz and Cdc42 resulted in supernumerary polar cells and was consequently associated with an increase in stalk cell number (Fig. 5L-N). Polar cell specific depletion of Dlg-1, however, did not affect polar cell number and yet significantly increased stalk cell numbers (Fig. 5L-N). Wildtype polar cells can be distinguished from surrounding main body follicle cells by strong expression of E-Cad (Fig. 5L, O) and display typical apical localization of Baz (Fig. 5O). Dlg-1 is strongly enriched between polar cell – main body follicle cell membranes but barely detectable at polar cell – polar cell membranes (Fig. 5O). While depletion of *cora* or *nrv2* did not affect E-Cad and Baz localization, polar cells failed to remove Dlg-1 from polar cell – polar cell membranes (Fig. 5P-R), indicating that SJ proteins are required in follicle cells to restrict Dlg-1 localization to appropriate membrane domains.

**Figure 5:**
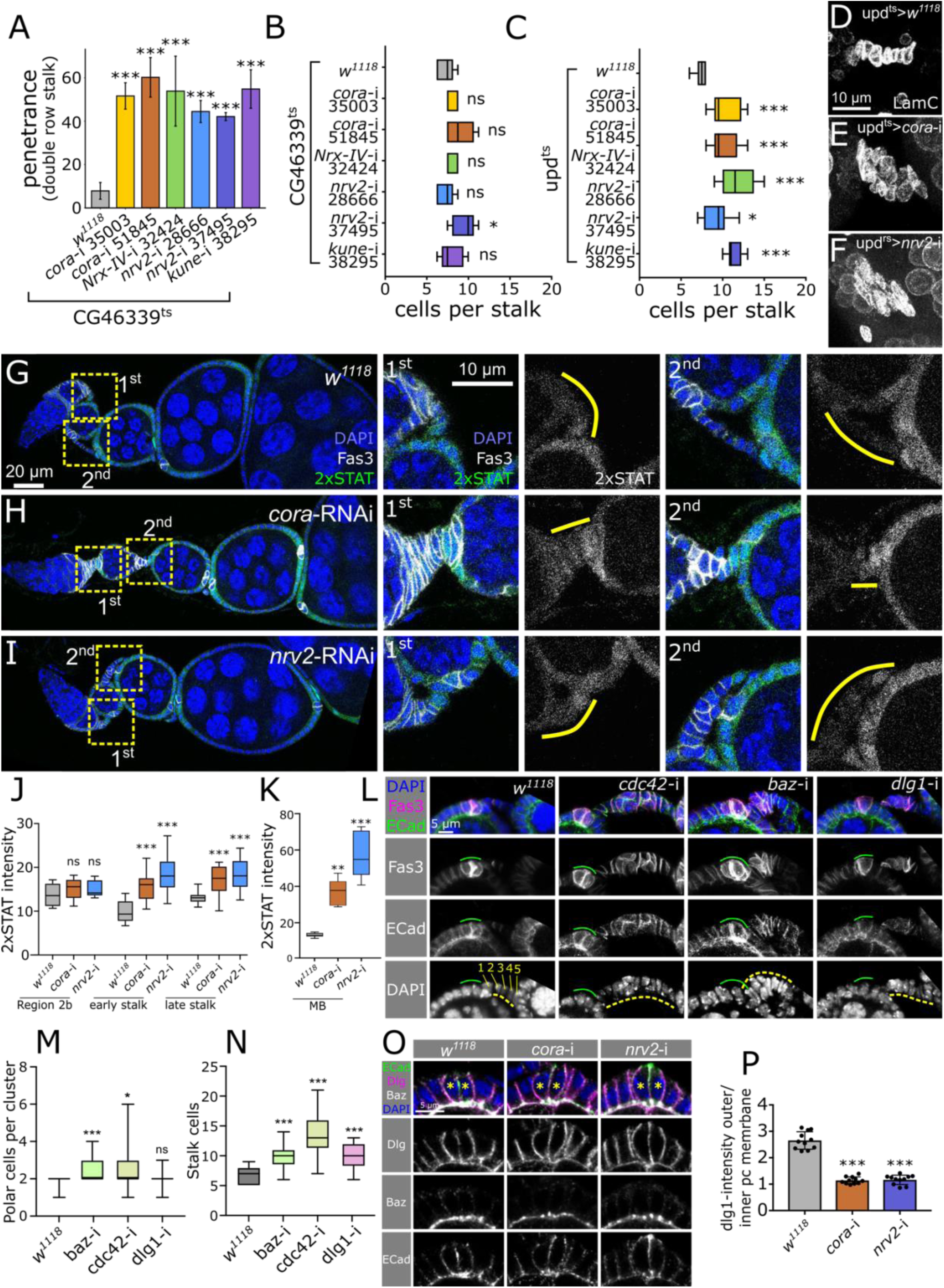
SJ proteins control polar cell polarity to regulate stalk cell specification. A) Quantification of the frequency of CG46339^ts^ ovarioles crossed to the respective allele with double row stalks (Stage 4 or older). P-values from Chi-squared test. n = 196, 150, 194, 180, 154, 135, 135 ovarioles respectively. B-C) Quantification of cells per Stage 4 stalk in ovarioles of CG46339^ts^ (B) or upd^ts^ (C) crossed to the respective genotype. Stalk cell specific knockdown of SJ components does not significantly increase stalk cell numbers or leads to only a mild increase (B) while polar cell specific depletion significantly increases stalk cell number (C). B) n = 9, 5, 5, 5, 5, 5, 5 stalks respectively, C) n = 8, 8, 8, 8, 10, 6 stalks respectively. D-F) Mature, posterior Stage 4 stalks stained for LamC in ovarioles with polar cell specific expression of the respective genotype. LamC is expressed in high levels in stalk cells and can be detected in lower levels in germline nuclei. Surrounding muscle cells also express LamC. G-I) Ovarioles of 109-30^ts^ combined with the Jak-STAT reporter UAS-2xSTAT-eGFP and the respective allele as indicated. We stained for DAPI (blue), GFP (green) and Fas3 (white) (G-I). Insets on the right show the first or second stalk as indicated and stained for DAPI (blue), GFP (green) and Fas3 (white) or 2xSTAT-GFP (white). J) Quantification of 2xSTAT-GFP in Region 2b, early stalk or mature stalk cells in 109-30^ts^ crossed to the respective genotype. n = 6, 5, 5, 14, 14, 14, 14, 14, 14 ovarioles respectively. K) Quantification of 2xSTAT intensity in the three main body follicle cells neighboring anterior polar cells in Stage 5-6 follicles in the background of 109-30^ts^ driving the indicated genotype. n = 5 ovarioles for each condition. L) Immunostaining of polar and stalk cells of Stage 5-6 ovarioles of 109-30^ts^ inducing the respective genotype and stained for DAPI, Fas3 and ECad. Fas3 is highly expressed in and ECad is enriched in polar cells. M-N) Quantifications of polar cells (M) and stalk cells (N) of Stage 5-6 of genotypes shown in L). Depletion of *baz* and *cdc42* increases polar and stalk cell numbers. *dlg-1* depletion does not affect polar cell number but significantly increases stalk cell number. M) n = 31, 40, 23, 33 polar cell clusters respectively. N) n = 15, 25, 20, 28 stalks respectively. O) Stage 6 polar cells of 109-30ts inducing the respective genotype and stained for DAPI, Dlg, Baz and ECad. Polar cells can be identified by enrichment of ECad in contrast to surrounding main body follicle cells. In control polar cells Dlg is enriched at polar cell – MB cell membranes and less present at polar cell – polar cell membranes. Polar cells with *cora* or *nrv2* depletion display a redistribution of Dlg while apical Baz localization is not affected. P) Quantification of Dlg fluorescence intensity at outer polar cell – MB cell membranes to inner polar cell – polar cell membranes. n = 10 measurements for each condition.

### Apical Upd enrichment is regulated by Cabeza-mediated mRNA transport

A previous study showed that *upd1*-mRNA is apically localized in polar cells (Ghiglione *et al*, 2002). Stalk cells face the basal side of polar cells, hence disturbed polar cell polarity may increase the availability of Upd1 ligand for stalk cells. We tested whether Upd1 protein is apically enriched in polar cells and indeed HA-tagged Upd1 primarily localized to the apical side of polar cells across all stages (Fig. 6A). We then tested the localization of Upd1 protein upon knockdown of *cora* and *nrv2* and found that Upd1 apical enrichment was abolished (Fig 6B-C). Simultaneously, knockdown of SJ components also resulted in a significantly higher recruitment of Upd1 protein to stalk cells (Fig. 6B, D). mRNA localization is normally mediated by motor protein driven transport along microtubules (Das *et al*, 2021). Using a fluorescent reporter for the microtubule plus-end binding protein Eb1, we found that, while wildtype polar cells accumulated Eb1 at the apical part of polar cells, *nrv-2* depletion disturbed apical enrichment of Eb1 (Fig. 6E-F). We tested for binding motifs of known RNA-binding factors in *upd1*-mRNA using BRIO (Guarracino *et al*, 2021) and identified two motifs for PUM2, the mammalian ortholog of *Drosophila* Pumilio (Pum) in the coding sequence of *upd1*-mRNA (Fig. S6A). Further, we identified three motifs for Fused in Sarcoma (FUS), orthologous to Cabeza (Caz), out of which one motif was localized in the coding sequence and two were found in the 3’ untranslated region (Fig. S6B). While Pum is well known for regulating mRNA stability, is was recently associated with mRNA transport in the nervous system (Grzejda *et al*, 2025; Nishanth & Simon, 2020). Caz, known for its functions in the nervous system (Sasayama *et al*, 2012; Xia *et al*, 2012; Shahidullah *et al*, 2013; Frickenhaus *et al*, 2015; Machamer *et al*, 2014) and contains an RNA recognition motif domain. Its mammalian ortholog FUS is associated with amyotrophic lateral sclerosis and has been shown to exhibit interaction with the plus-end directed motor protein Kinesin (Yasuda *et al*, 2017). Depletion of either *pum* or *caz* from follicel cells resulted in defective stalk morphogenesis (Fig. 6G-H), prompting whether apical localization of Upd depended on either factor. While Upd-HA primarily localized to the apical part in *pum* depleted polar cells, similar to the wildtype control, depletion of *caz* disrupted Upd apical localization entirely (Fig. 6I).

**Figure 6:**
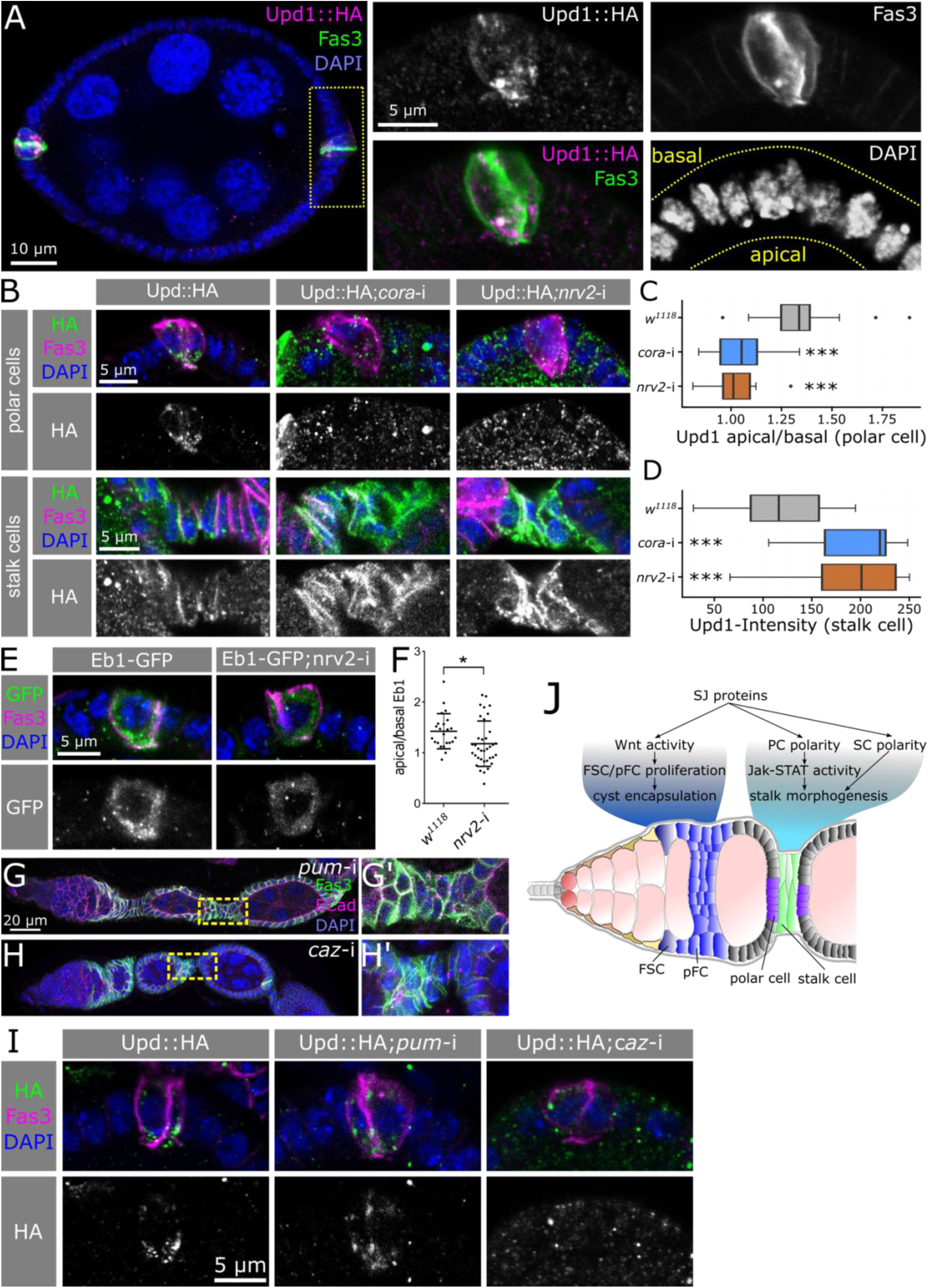
Apical enrichment of Upd ligand relies on SJ components. A) A Stage 6 cyst expressing Upd1::HA stained for HA (magenta), Fas3 (green) and DAPI (blue) and insets showing the indicated channels Yellow outline in the DAPI channel marks the epithelium. B) Stage 4-6 polar cells (upper panels) and stalks (lower panels) of upd^ts^ inducing Upd::HA expression and the indicated RNAi, if applicable. We stained for HA (green or white) to visualize Upd localization, DAPI (blue) for nuclei and Fas3 (magenta), which is highly expressed in polar cells. In wildtype polar cells, Upd is enriched at the apical polar cell side. This enrichment is disturbed upon depletion of *cora* or *nrv2*. SJ component depletion also results in higher Upd signal in stalk cells. C-D) Quantification Upd intensity in genotypes shown in B). C) Quantification of apical versus basal Upd in polar cells. n = 20 polar cells for each condition. D) Quantification of Upd in stalk cells. n = 18, 17, 19 stalks respectively. E-F) Stage 6 polar cells expressing Eb1-GFP in the 109-30ts background or with additional *nrv2*-RNAi stained for GFP (green/white), Fas3 (magenta) and DAPI (blue) and associated quantification of Eb1 intensity in apical versus basal polar cell parts (F). G-H) Ovarioles of 109-30^ts^ driving RNAi targeting *pum* (G) or *caz* (H) stained for Fas3 (green), ECad (magenta) and DAPI (blue). G’-H’) show insets at stalk regions. I) Stage 4-6 polar cells of 109-30^ts^ driving Upd::HA in the control background or combined with RNAi targeting *pum* or *caz*. J) Graph summarizing the functions of SJ core components in early follicle cell development. SJs allow normal FSC/pFC proliferation rates by promoting Wnt signaling activity which is vital for effective cyst encapsulation in Region 2b. SJs further regulate stalk morphogenesis non-cell-autonomously by maintaining cell polarity in polar cells, allowing Caz-mediated apical transport of *upd1*-mRNA and thereby fine-tuning Jak-STAT signaling activity. Additionally, they cell-autonomously control stalk cell apical-basal polarity, cell adhesion and actin cytoskeleton to coordinate the intercalation process.

## Discussion

In this study we identified several functions for Cora, Nrv2, Nrx-IV and Kune during follicle epithelial development unrelated to the occluding function of SJs (Fig. 6J). By regulating cell polarity, these SJ proteins facilitate Wnt signaling in undifferentiated follicle cells to ensure normal proliferation levels, which is the prerequisite for effective germline cyst encapsulation. They further regulate polar cell polarity allowing caz-mediated apical transport of *upd1*-mRNA and thereby fine-tune Jak-STAT signaling levels in surrounding follicle cells. Lastly, SJ proteins are vital for normal stalk morphogenesis by maintaining apical-basal polarity and governing F-actin enrichment on the apical side, which provide driving forces for the stalk cell intercalation process. Together, our study demonstrates that SJ components and normal apical-basal polarity are indispensable for normal epithelial development from the stem cell onwards.

### SJ proteins mediated cell polarity governs follicle epithelial development

Mechanistically, we find that SJ proteins function via the regulation of apical-basal polarity. Cora, Nrv2 and Nrx-IV have previously been associated with cell polarity regulation during embryonic epithelial development (Laprise *et al*, 2009) and we identify Kune as an additional SJ protein involved in polarity regulation. SJ proteins localize inter-dependently in follicle cells, suggesting that each of the tested components function in the same pathway. As mature SJs are formed well beyond the Stages we analyzed (Müller, 2000), our data provide solid evidence that cell polarity control is exerted by SJ components in a non-occluding manner. SJ protein or Dlg-1 depletion diminishes Wnt pathway activity in FSCs alike. Importantly, FSCs do not contain a defined apical membrane domain. While Dlg-1 is found at FSC membranes, Baz is the first apical determinant localizing apically from the pFC state onwards, where it is involved in positioning adherens junctions and inducing apical-basal polarity (Kronen *et al*, 2014; Franz & Riechmann, 2010). As both Dlg-1 as well as SJ components are required for appropriate Wnt signaling activity, these proteins may assume key regulator roles during follicle cell polarity establishment. In stalk and polar cells, SJ proteins are vital to restrict Dlg-1 from appropriate membrane compartments suggesting a similar mechanism in both cell types. Contrastingly, studies on embryonic epithelial polarity found no genetic interaction between SJ proteins and Dlg-1 and instead proposed that SJ proteins function opposingly to the apical Crumbs complex (Laprise *et al*, 2009). In the future, further studies are required to dissect the mechanistical differences of embryonic and follicle epithelial cell polarity establishment.

### SJ proteins regulate proliferation via the Wnt signaling pathway

SJ proteins have been linked to proliferation in the midgut epithelium, where they limit intestinal stem cell proliferation, and in the imaginal disc epithelium, where *cora* promotes proliferation (Chen *et al*, 2020; Izumi *et al*, 2019; Ward *et al*, 2001). We found that SJ components are required for sustained FSC and pFC proliferation but are dispensable for main body follicle cell proliferation. This highlights that regulation of proliferation by SJ components is highly tissue specific. We identify a previously unrecognized function of SJ components in promoting Wnt signaling to induce proliferation. Depletion of SJ genes or positive regulators of Wnt signaling similarly results in an accumulation of cysts in Region 2b, where germline cysts are first encapsulated by follicle cells. Normal levels of proliferation, regulated by SJ proteins through Wnt signaling activity, thus are necessary to ensure the production of sufficient pFCs to populate entering germline cysts. Our studies find that Dlg-1 alike is required for promote Wnt signaling, suggesting that Wnt pathway activity is regulated by cell polarity. Frizzled proteins, the receptors for the Wnt family of signaling ligands, display apical-basal polarization in *Drosophila* as well as mammals (Wang *et al*, 2006; Wu *et al*, 2004). It is conceivable, that the one or several Frizzled receptors are distinctly localized by SJ proteins and Dlg-1 in the FSC to calibrate Wnt activity. However, due to a limitation of functional antibodies and reporter lines, we were unable to test this hypothesis. As FSCs do not exhibit mature apical-basal polarity (Kronen *et al*, 2014; Franz & Riechmann, 2010), other alternative hypothesis are possible. SJs may be required for FSC adhesion thereby regulating exposure to Wnt ligands provided by niche cells. One apparent candidate is E-Cad, which is crucial for FSC niche adhesion (Song & Xie, 2002) and is regulated by SJ proteins in stalk cells. However, we did not observe any obvious changes in E-Cad localization or expression in Region 2b follicle cells upon SJ component knockdown (data not shown). A previous study has suggested a role for the SJ protein Neuroglian in lateral cell adhesion and SJ proteins likely regulate cell-cell adhesion during border cell migration in Stages 9 to 10 egg chambers (Alhadyian *et al*, 2021; Bergstralh *et al*, 2015). These findings point to the exciting possibility that SJ proteins may be directly required for FSC niche adhesion.

### Cell polarity and the actin cytoskeleton are required for stalk cell intercalation

SJ components regulate stalk cells in a cell autonomous way. Stalk cell morphogenesis is a poorly understood process. While it is known that the basement membrane is important for stalk cell morphogenesis, cell polarity and cytoskeleton dynamics have not been investigated (Van De Bor *et al*, 2021). Here, we describe the distinct steps of stalk cell morphogenesis and provide evidence that E-Cad-mediated cell adhesion and the actin cytoskeleton are key drivers of stalk intercalation. Using *ex vivo* live imaging, we further show that stalk cell intercalation occurs over the course of a few hours. SJ proteins are required for normal localization of E-Cad and of the polarity proteins Baz and Dlg-1. Moreover, SJ proteins are vital for apical accumulation of F-actin in intercalating stalk cells. SJs have also been shown to regulate E-Cad and actomyosin during dorsal closure in *Drosophila* embryonic development and mediate actin dynamics during wound closure (Carvalho *et al*, 2018; De *et al*, 2022). Together, these data support a central role of SJ proteins in dynamic processes dependent on cellular reorganization.

### SJ proteins are vital of apical Upd localization

SJ proteins further regulate stalk development non-cell autonomously by affecting Upd ligand provision from polar cells, which promotes stalk differentiation via Jak-STAT signaling (Assa-Kunik *et al*, 2007; McGregor *et al*, 2002). SJ proteins maintain normal localization of Dlg-1 and microtubule orientation in polar cells, a prerequisite of apical *upd1*-mRNA localization. We identified Caz, whose ortholog FUS plays critical roles in amyloid lateral sclerosis (ALS), as a RNA-binding protein with several recognition motifs in the *upd1*-mRNA 3’UTR. Our data, for the first time, suggest a role for Caz in RNA-transport. During ALS, FUS forms inclusions, in which it sequesters both Kinesin protein as well as mRNA (Yasuda *et al*, 2017), making it tempting to speculate that Caz can similarly bind to Kinesin to promote plus-end directed transport of *upd1*-mRNA towards the apical side of polar cells. The canonical function of FUS includes the regulation of DNA damage repair, a process that also relies on nuclear microtubules (Sama *et al*, 2014; Oshidari *et al*, 2018). A growing body of evidence further links RNA biology to DNA damage repair (Bader *et al*, 2020). It remains to be seen whether FUS could act as a linker between these processes.

Apical enrichment of Upd ligand, regulated indirectly by SJ proteins, is vital to limit Jak-STAT activity in stalk cells. SJ proteins also regulate egg chamber elongation during late stages of oogenesis although the underlying mechanism is not understood (Alhadyian *et al*, 2021). Importantly, egg chamber elongation is dependent on Jak-STAT signaling induced by polar cells and overexpression of Upd results in failed elongation (Alégot *et al*, 2018). Our findings indicate that SJ proteins regulate egg chamber elongation by limiting Jak-STAT pathway activity in main body follicle cells.

Interestingly, in the midgut epithelium smooth SJ members directly interact with the Hippo signaling effector Yorkie to limit Upd ligand expression (Chen *et al*, 2020; Izumi *et al*, 2019). In polar cells, forced nuclear entry of Yorkie results in the loss of polar cells and subsequently stalk cells (Chen *et al*, 2011). Since SJ component depletion does not alter polar cell numbers, they arguably do not regulate Jak-STAT activity via Hippo signaling in the follicle epithelium. Instead, they regulate aspects of cell polarity in polar cells, which can contribute to the apical localization and thereby distribution of Upd1. Fascinatingly, our study and others therefore show that SJ components regulate the Jak-STAT signaling pathway in distinct, cell-type specific ways (Chen *et al*, 2020; Izumi *et al*, 2019).

### Conserved non-barrier functions of occluding junctions

SJ proteins, including Cora, Nrv2, Nrx-IV and Kune, are conserved in vertebrates, where their homologs contribute to axo-glial SJs (Banerjee *et al*, 2006; Faivre-Sarrailh, 2020). These vertebrate SJs, similar to their invertebrate counterpart, form a diffusion barrier for paracellular transport and limit lipid exchange along the cell membrane. Consistently, mutations in axo-glial SJ genes are associated with a range of different neuropathies (Faivre-Sarrailh, 2020). While non-occluding functions of axo-glial SJs have not yet been described, tight junctions, the vertebrate epithelial occluding junctions, fulfill several non-barrier functions. Like SJs, they reciprocally interact with the apical-basal polarity machinery (Shin *et al*, 2006). Data from our study and others show that this function is conserved in invertebrate occluding junctions (this study and (Laprise *et al*, 2009)). Tight junctions have also been linked to cell cycle regulation but a recent review highlights that the interplay with proliferation regulating signaling pathways is insufficiently understood (Nehme *et al*, 2023). In this study we identify a novel link between SJ components and the Wnt and Jak-STAT signaling pathways. Other research has established that SJ components further regulate Hippo signaling (Khadilkar & Tanentzapf, 2019; Xu *et al*, 2019). It is plausible that vertebrate occluding junctions similarly regulate a variety of signaling pathways that coordinate proliferation and differentiation. This association is of particular interest in the background of a growing body of evidence that connects tight junctions to cancer development (Nehme *et al*, 2023).

## Material and Methods

### Fly husbandry and fly stocks

Flies were reared under standard lab conditions. All crosses including a tub-Gal80^ts^ construct were reared at 18°C and hatched offspring were transferred to 29°C for 14 days and fed wet yeast daily until dissection.

The following stocks were used in this study:

BDSC: *w^1118^* (3605), 109-30-Gal4 (7023), CG46339-Gal4 (77710), tub-Gal80ts (7108), UAS-*cora*-RNAi (35003), UAS-*cora*-RNAi (51845), UAS-*kune*-RNAi (38295), UAS-*nrv2*-RNAi (28666), UAS-*nrv2*-RNAi (37495), UAS-*Nrx-IV*-RNAi (32424), UAS-armS10 (4782), UAS-*baz*-RNAi (35002), UAS-*shg*-RNAi (32904), UAS-*pum*-RNAi (38247), UAS-2XStat92E-GFP (26196), Nrx-IV-mCherry (92348). VDRC: UAS-*arm*-RNAi, (7767), UAS-*dsh*-RNAi (101525), UAS-*dlg1*-RNAi (41134), UAS-*cdc42*-RNAi (100794), UAS-*caz*-RNAi (100291). upd-Gal4 was kindly gifted from Denise Montell. fz3::RFP was a kind gift from Ramanuj DasGupta. upd::HA was kindly shared by Hiroshi Nakato. ECad-3xGFP was kindly provided by Jörg Grosshans. We used BDSC #51845 for *cora* knockdown and BDSC #28666 for *nrv2* knockdown if not specified otherwise.

### Immunofluorescence staining and imaging

Protocols were carried out at room temperature when not indicated otherwise. Flies were dissected in PBS and ovaries fixed for 20 minutes in 4% PFA. Ovaries were washed thrice for 10 minutes with PBS and blocked for 1h with block solution (PBS with 0,2% Triton X-100 and 0,5% BSA). Primary antibody incubation was performed overnight at 4°C in block solution. For F-actin staining phalloidin (Invitrogen A12380, 1:200) was added to the solution. On the next day the ovaries were washed three times for 10 minutes with block solution and incubated with secondary antibody in block solution for 2-3h. Lastly, ovaries were washed three times for 10 minutes with block solution and twice with PBS. Ovaries were mounted in DAPI Fluoromount-G (ThermoFisher Scientific OB010020).

The following antibodies were used:

DSHB: mouse anti-Cora (1:1000, C566.9), mouse anti-Hts (1:100, 1B1), mouse anti-Fas3 (1:100, 7G10), mouse anti-LamC (1:100, LC28.26), mouse anti-Yan (1:100, 8B12H9), mouse anti-Eya (1:100, 10H6), rat anti-E-Cad (1:50, DCAD2), mouse anti-Dlg1 (1:50, 4F3). Cell Signaling: rabbit anti-c-Dcp1 Asp215 (1:100, #9578), rabbit anti-GFP (1:1000, D5.1). Rabbit anti-HA (1:100, Invitrogen SG77), rat anti-RFP (1:1000, ChromoTek 5F8). Rabbit anti-Baz (1:100, kind gift from Andreas Wodarz), guinea pig anti-zfh1 (1:500, kind gift from James Skeath). Secondary antibodies were purchased from Invitrogen and used at 1:1000: goat anti-guinea pig 488 (A11073), goat anti-mouse 488 (A11029), goat anti-mouse 568 (A11031), goat anti-mouse 647 (A21230), goat anti-rabbit 488 (A11034), goat anti-rabbit 568 (A11036), goat anti-rat 488 (A11006).

For the EdU assay ThermoFisher EdU Alexa Fluor 555 Imaging Kit (C10338) was used. Ovaries were dissected in PBS and incubated in 15 µm EdU in PBS for 1h. Ovaries were washed two times with PBS and fixed for 20 minutes in 4% PFA. Ovaries were blocked for 1h in blocking solution and primary antibody was incubated overnight at 4°C. Ovaries were washed twice with PBS and incubated 30 minutes with Click-iT reaction cocktail and covered. Ovaries were washed two times with PBS and blocked for 1h. Secondary antibody was incubated overnight at 4°C. Ovaries were washed thrice with blocking solution and twice with PBS and mounted in DAPI Fluoromount-G.

Images were acquired with Leica TCS SP8 with HC PL APO CS2 40x/1.30 and HC PL APO CS2 x63/1.4 oil objective and Leica Application Suite X (LasX) software. For intensity measurements images were acquired with the same settings. Image processing, analysis and fluorescence intensity measurements were performed with FIJI 2.9.0 (Schindelin et al., 2012). Dlg1 and Actin colocalization was assessed along the zipper line with the JACoP tool v2.0 from the BIOP plugin in FIJI. Wnt signaling activity was assessed by measuring the fluorescence intensity of fz3::RFP. To control our results between specimen, we normalized each measurement point in Region 2b with a measurement in Region 1, where cells are not affected by our Gal4-driver 109-30. Figures were prepared using Inkscape 1.3.2.

EdU positive cell quantification was performed with Imaris 9.3 (Bitplane). Images were imported as z-projections and a region of interest was selected. The total cell count within this region was determined using the DAPI channel spot tool. EdU positive cells were counted within the same region using the RFP channel. Stalk cross-sections were visualized with Imaris 9.3 (Bitplane) using the E-Cad channel of z-stacks taken with 0.75 µm distance.

### Live imaging of *Drosophila* ovarioles

Flies were wet yeasted at for least 3 days at 25°C and ovaries dissected in Scheider^++^ medium supplemented with 20 µg/ml bovine insulin. Young stages were embedded in Scheider^++^ medium containing 20 µg/ml bovine insulin and 0.5% low melting agarose in an 8-well imaging chamber (ibidi). Imaging was performed with a Zeiss CellObserver Z.1 with a Yokogawa CSU-X1 spinning disk scanning unit and an Axiocam MRm CCD camera.

### Statistics and reproducibility

Expression of SJ components was assessed using a publicly available single cell RNA-sequencing atlas (Rust et al., 2020) via the DotPlot() function of the Seurat package (5.0.2) in RStudio (2023.12.1+402).

Significance tests were performed with Prism (10) or RStudio (2024.04.0+735). We tested normal distribution with the Shapiro-Wilk test. For normally distributed samples we then performed a student’s t-test. For all other samples we performed the Mann-Whitney test. For significance tests of more than two groups we tested for normality and performed the Kruskal-Wallis test. Pearson’s Chi-squared test with Yates correction was performed with the chisq.test() command of the stats package (4.3.2) in RStudio (2024.04.0+735). Significance values are reported as: ns > 0.05, * < 0.05, ** < 0.01, *** < 0.001. In box plots, the midline depicts the median, the lower and upper hinges correspond to the 25 and 75 percent quartiles and the whiskers span the smallest and largest values. Quantifications in the main text are reported as arithmetic mean ± SEM. All images and quantifications are representatives of at least 3 technical and 15 biological replicates.

## Data availability

The data that support the findings of this study are available within the article, Supplementary Information, or from the corresponding author upon request. Source data are provided with this paper.

## Acknowledgements

We thank the Bloomington and Vienna Stock centers and Denise Montell and Jörg Grosshans for fly stocks and Andreas Wodarz and James Skeath for kindly gifted antibodies. We are grateful to Sona Martirosan for the initial assessment of phenotypes caused by stalk and polar cell specific knockdown of SJ components. We thank Andreas Wodarz, Sven Bogdan and Dennis Klug for constructive comments on the manuscript. This work was supported by the PSI (Promoting Scientific Independence) Program of the Philipps-University Marburg. This work was supported by the Deutsche Forschungsgemeinschaft (DFG, German Research Foundation) – Project number 558245783.

## Author contributions

K.R. designed the project and wrote the manuscript. V.H. and K.R. made the figures. A.S, V.H. and H.M performed dissections and immunofluorescence stainings. V.H. performed microscopy, quantifications and statistical analyses. All authors commented on the manuscript.

## Competing interests

The authors declare no competing interests.

---

**Supplementary Figure 1:**
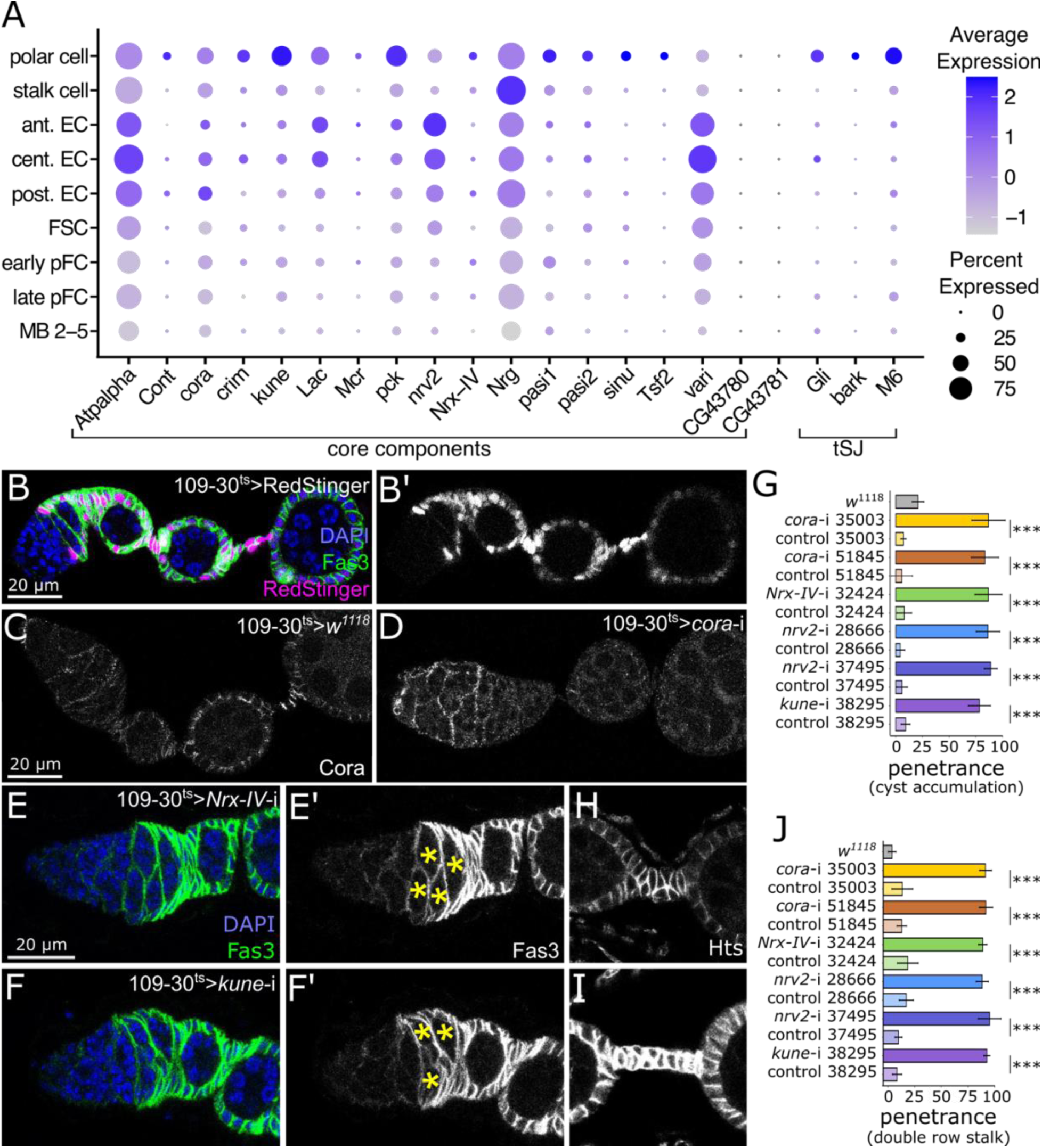
SJ components are expressed in and required for follicle cells. a) Average and percent expression of SJ core components and tricellular SJ (tSJ) components in escort cells and early follicle cell types identified in (Rust *et al*, 2020). Many SJ components display high expression in polar cells and several components are enriched in escort cells. B) Ovariole expressing RedStinger (magenta) under the 109-30-Gal4 driver and stained for Fas3 (green) and DAPI (blue). B’) RedStinger expression in white. C-D) Ovarioles of 109-30^ts^ crossed to *w^1118^* (C) or *cora*-RNAi (D) and stained for Cora. E-F) Ovarioles with 109-30^ts^ induced depletion of *Nrx-IV* (E) or *kune* (F) stained for DAPI (blue) and Fas3 (green). G) Quantification of the frequency of ovarioles with more than two germline cysts in Region 2b of 109-30^ts^ crossed to the respective allele. For control genotypes the CyO positive sibling flies, which lack the 109-30-Gal4 allele, were dissected. P-values from Chi-squared test. n = 247, 236, 184, 219, 154, 232, 161, 252, 153, 150, 134, 158, 142 germaria respectively. H-I) Stalk regions of ovarioles of 109-30^ts^ crossed to *Nrx-IV*-RNAi (H) or *kune*-RNAi (I) stained for Hts. J) Quantification of the frequency of ovarioles with double row stalk (Stage 4 or later) of 109-30^ts^ crossed to the respective allele. Control flies were devoid of 109-30-Gal4. P-values from Chi-squared test. n = 197, 138, 149, 140, 135, 140, 136, 148, 144, 142, 133, 140, 147 ovarioles respectively.

**Supplementary Figure 2:**
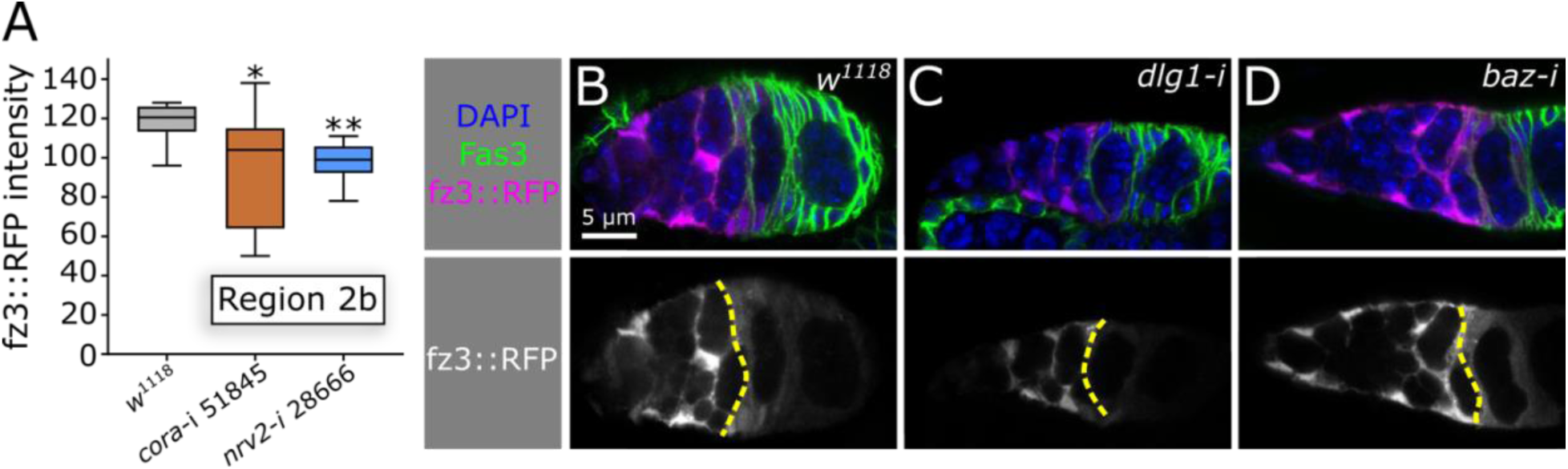
Cell polarity affects Wnt signaling activity in early follicle cells. A) Quantification of fz3::RFP intensity in Region 2b for 109-30^ts^ inducing each respective genotype n = 8, 9, 6 germaria respectively. B-D) Ovarioles of 109-30ts combined with fz3::RFP (magenta) and the respective genotype stained for Fas3 (green) and DAPI DAPI). Lower panel shows the fz3::RFP in white. Yellow dotted line marks the Fas3 boundary.

**Supplementary Figure 3:**
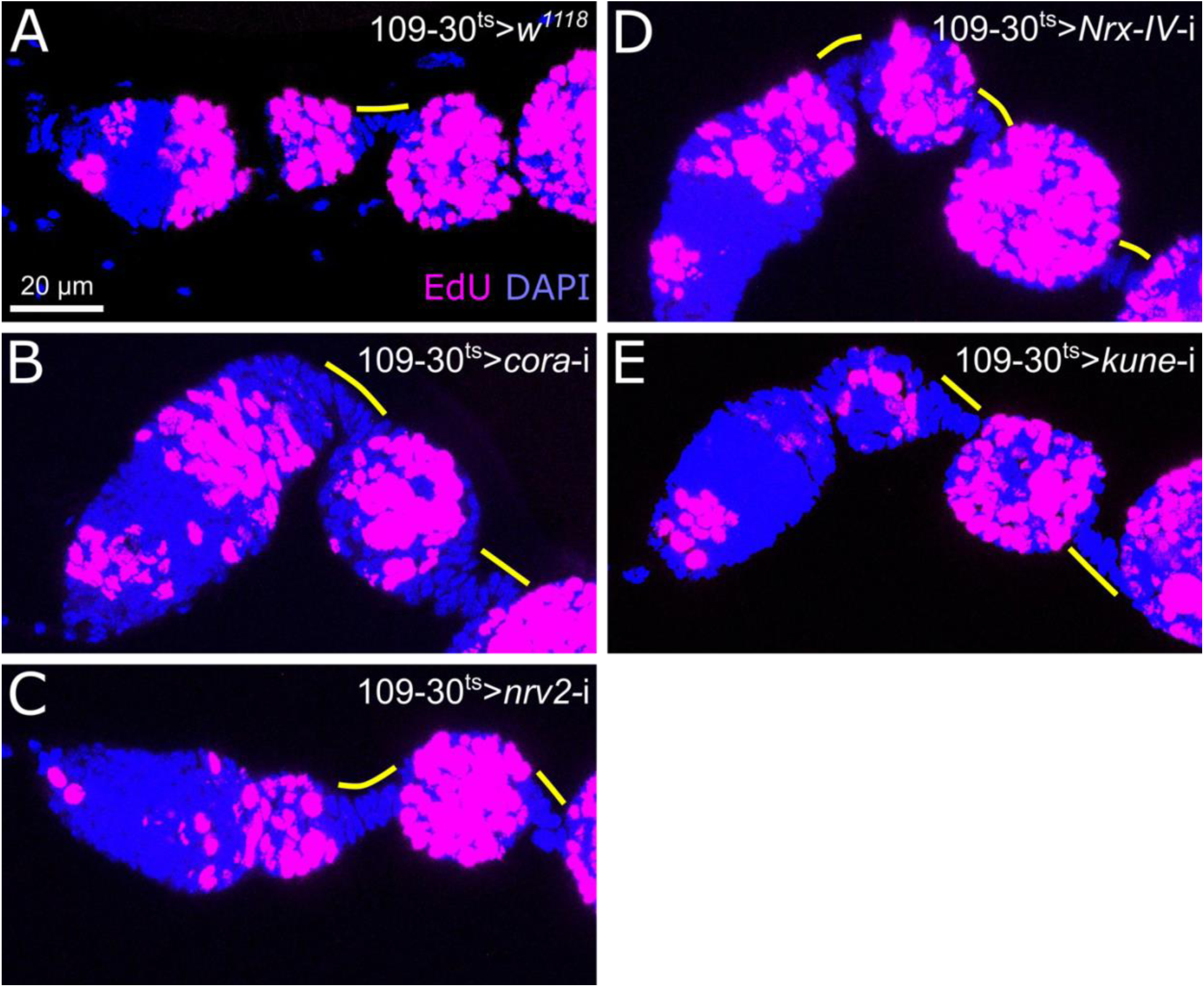
SJ component depletion does not promote stalk cell proliferation. A-E) Ovarioles of the respective genotypes stained for DAPI (blue) and EdU (magenta). Stalk cells are EdU negative and outlined in yellow.

**Supplementary Figure 4:**
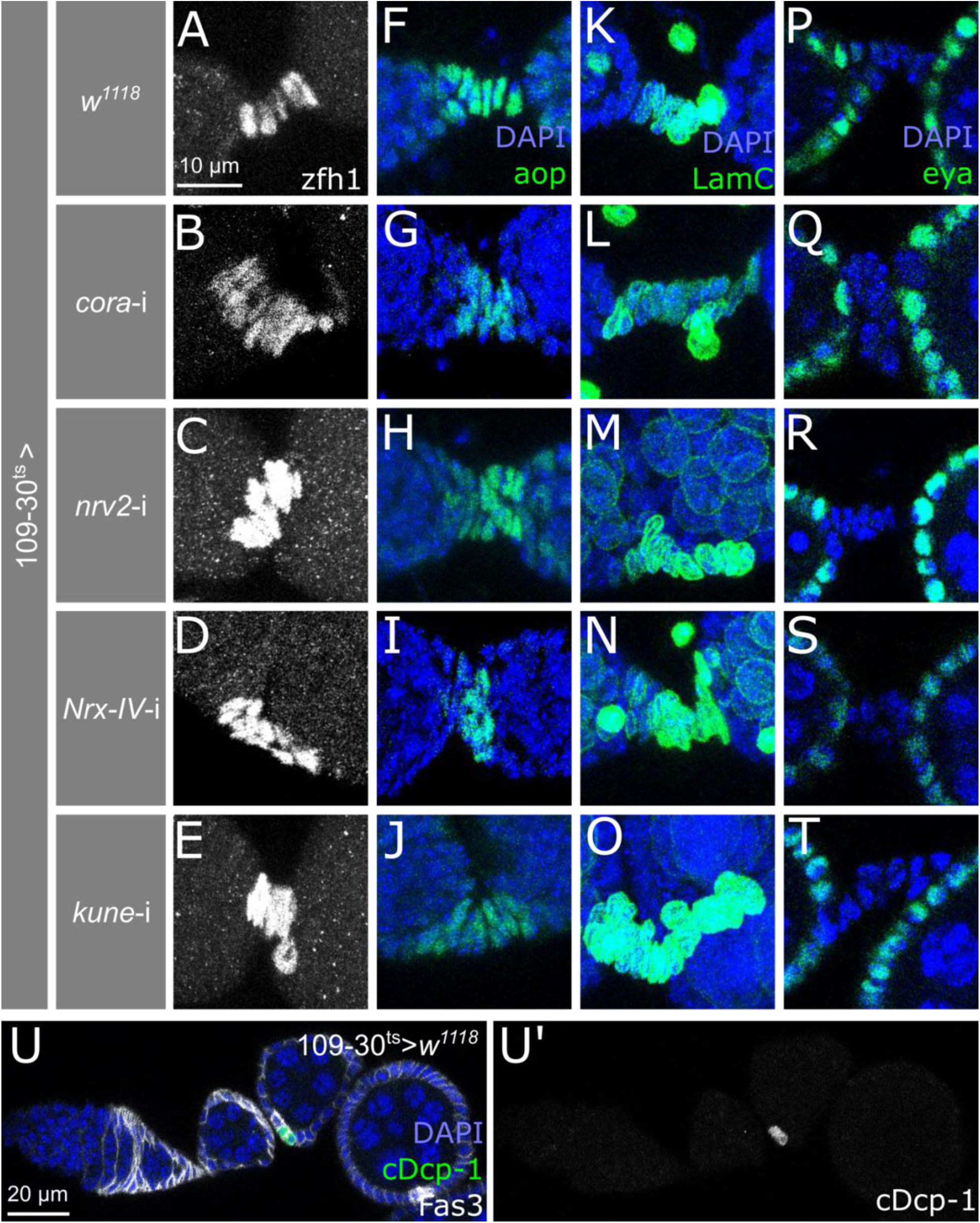
SJ proteins do not affect stalk cell differentiation or apoptosis. A-T) Stalk regions of 109-30ts inducing the respective genotype stained for cell type specific markers. A-E) zfh1 (white) is expressed in stalk cells. F-O) aop and LamC (green) are expressed in a stalk cell specific manner. P) eya (green) is excluded from stalk cells. F-O) DAPI staining in blue. U) 109-30^ts^ > *w^1118^* ovariole stained for DAPI (blue), c-Dcp-1 (green) and Fas3 (white). U’) shows c-Dcp-1 (white). We detected c-Dcp-1 staining in Fas3 high polar cells but not in stalk cells.

**Supplementary Figure 5:**
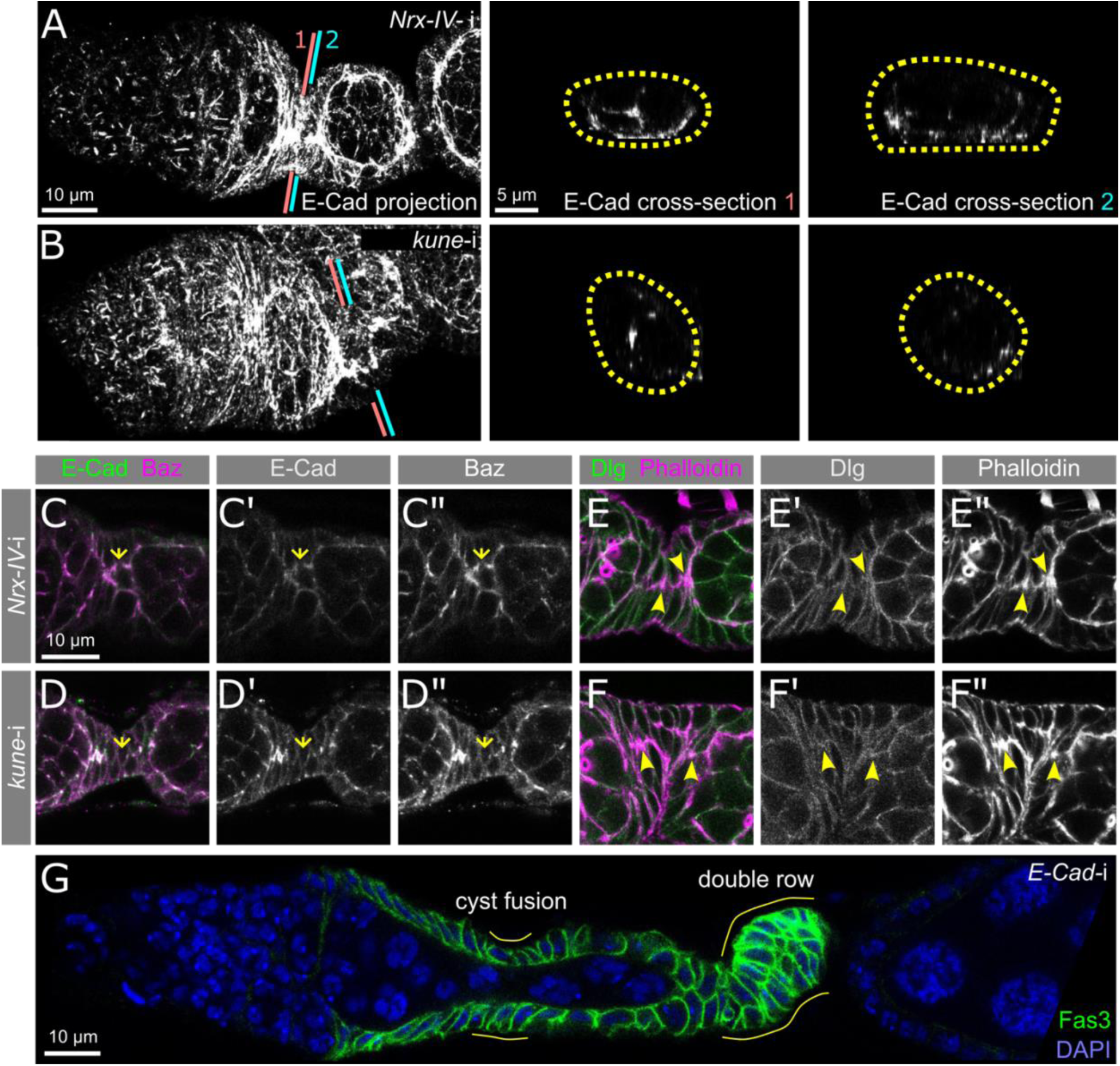
SJ proteins and E-Cad are required for stalk cell intercalation. A-B) Maximum intensity projections and stalk cross sections of E-Cad staining in 109-30^ts^ driving *Nrx-IV*-RNAi (A) or *kune*-RNAi (B). C-F) Immunostainings of budding stalk regions of 109-30^ts^ driving *Nrx-IV*-RNAi (C, E) or *kune*-RNAi (D, F) stained for E-Cad and Baz (C-D) or Dlg and Phalloidin (E-F) as indicated. G) Ovariole with pan-follicle cell depletion of *E-Cad* (109-30^ts^ driving *E-Cad*-RNAi) stained for Fas3 (green) and DAPI (blue). Areas with cyst fusion and double row stalks are outlined in yellow.

**Supplementary Figure 6:**
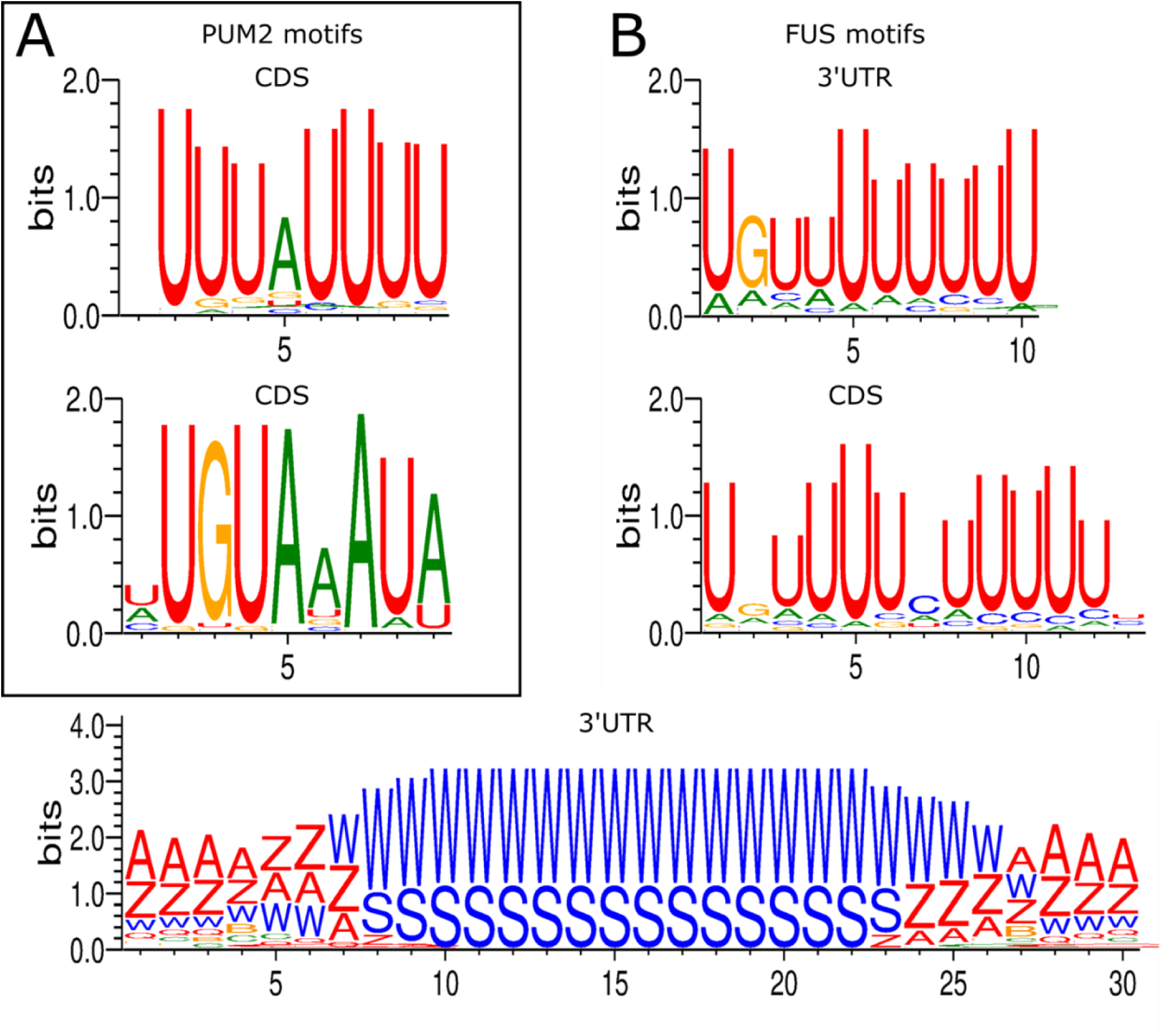
Recognition motifs for RNA-binding proteins in upd1-mRNA. A-B) BRIO analysis identified binding motifs for PUM2 (orthologous to *pum*) and FUS (orthologous to *caz*) in *upd1*-mRNA. A) PUM2 recognition motifs are found in the coding sequence (CDS). P-values: 0,031 (upper panel), 0,037 (lower panel). B) FUS motifs are found in the 3’ untranslated region (UTR) and the coding sequence (CDS). The upper-most motif is present 14 times in the *upd1* 3’UTR with a p-value of 0,014. Middle panel: p = 0,039, lowest panel: p = 0,042.

## Notes

### Competing Interest Statement

The authors have declared no competing interest.

### Summary of Updates

The updated version contains new data confirming that SJ proteins regulate cell polarity throughout follicle epithelial development. We confirm that SJ proteins and cell polarity affect Wnt signaling in undifferentiated follicle cells. Further, we identify a previously unknown process of Jak-STAT signaling regulation. SJ proteins regulate cell polarity and microtubule orientation in ligand-producing cells. This is a prerequesite for apical transport of upd1-mRNA via the RNA-binding protein Cabeza, the ortholog of which (FUS) is known to interact with kinesin motor protein. The apical enrichment of upd1-mRNA and subsequently Upd protein limits the efficiency with which basally located stalk cells are specified.

